# CSN and CAVA: variant annotation tools for rapid, robust next-generation sequencing analysis in the clinic

**DOI:** 10.1101/016808

**Authors:** Márton Münz, Elise Ruark, Anthony Renwick, Emma Ramsay, Matthew Clarke, Shazia Mahamdallie, Victoria Cloke, Sheila Seal, Ann Strydom, Gerton Lunter, Nazneen Rahman

**Affiliations:** Wellcome Trust Center for Human Genetics, Roosevelt Drive, Oxford, OX3 7BN, UK; Division of Genetics & Epidemiology, Institute of Cancer Research, 15 Cotswold Road, London, SM2 5NG, UK; TGLclinical, Institute of Cancer Research,, 15 Cotswold Road, London, SM2 5NG, UK; The Royal Marsden NHS Foundation Trust, Downs Road Sutton, SM2 5PT, UK

**Keywords:** Clinical genomics, next-generation sequencing, variant annotation, indel, mutations

## Abstract

**Background:** Next-generation sequencing (NGS) offers unprecedented opportunities to expand clinical genomics. It also presents challenges with respect to integration with data from other sequencing methods and historical data. Provision of consistent, clinically applicable variant annotation of NGS data has proved difficult, particularly of indels, an important variant class in clinical genomics. Annotation in relation to a reference genome sequence, the DNA strand of coding transcripts and potential alternative variant representations has not been well addressed. Here we present tools that address these challenges to provide rapid, standardized, clinically appropriate annotation of NGS data in line with existing clinical standards.

**Methods:** We developed a clinical sequencing nomenclature (CSN), a fixed variant annotation consistent with the principles of the Human Genome Variation Society (HGVS) guidelines, optimized for automated variant annotation of NGS data. To deliver high-throughput CSN annotation we created CAVA (Clinical Annotation of VAriants), a fast, lightweight tool designed for easy incorporation into NGS pipelines. CAVA allows transcript specification, appropriately accommodates the strand of a gene transcript and flags variants with alternative annotations to facilitate clinical interpretation and comparison with other datasets. We evaluated CAVA in exome data and a clinical *BRCA1/BRCA2* gene testing pipeline.

**Results:** CAVA generated CSN calls for 10,313,034 variants in the ExAC database in 13.44 hours, and annotated the ICR1000 exome series in 6.5 hours. Evaluation of 731 different indels from a single individual revealed 92% had alternative representations in left aligned and right aligned data. Annotation of left aligned data, as performed by many annotation tools, would thus give clinically discrepant annotation for the 339 (46%) indels in genes transcribed from the forward DNA strand. By contrast, CAVA provides the correct clinical annotation for all indels. CAVA also flagged the 370 indels with alternative representations of a different functional class, which may profoundly influence clinical interpretation. CAVA annotation of 50 *BRCA1/BRCA2* gene mutations from a clinical pipeline gave 100% concordance with Sanger data; only 8/25 *BRCA2* mutations were correctly clinically annotated by other tools.

**Conclusions:** CAVA is a freely available tool that provides rapid, robust, high-throughput clinical annotation of NGS data, using a standardized Clinical Sequencing Nomenclature.

## BACKGROUND

Genetic testing has been an important clinical activity for over 20 years during which time many different mutation detection methods have been utilized and many thousands of clinically-relevant variant datasets have been generated. In recent years next-generation sequencing (NGS) has been transforming clinical genomics, allowing rapid interrogation of tens of thousands of genes and the identification of millions of variants [1]. Integration of pre-NGS and NGS data are essential for the correct interpretation and management of variants in the clinic, particularly as most clinical laboratories continue to use non-NGS methods for at least some tests (e.g. testing for individual mutations).

There are important, underappreciated differences in the outputs of pre-NGS and NGS gene sequencing methods which are hindering the required integration of data and thus the potential of genomics to impact health. The most pressing issue requiring attention is the huge variability in descriptive terminology of variants which is endemic both within and between pre-NGS and NGS annotation systems. For example, rs80357713 is the identifier of one of most well documented variants in the world; an Ashkenazim *BRCA1* founder mutation. Currently, rs80357713 is associated with 10 different annotations on dbSNP, none of which is the standard clinical representation of the mutation: *BRCA1* c.68_69delAG [2].

Clinical annotation of pre-NGS sequence data is generally in accordance with the Human Genome Variation Society (HGVS) guidelines [3]. However, these permit alternative annotations of some variants and hence foster inconsistency. They also allow terms that are incompatible with contemporary large-scale variant databases, such as an asterisk (which is used as a wildcard term in many applications) for stop-gain mutations. Currently, there are no universal annotation standards for describing NGS data, with different tools using similar, but not identical, notation systems [4-6]. A fixed, standardized, versioned nomenclature for clinical sequence data, identical for all mutation detection platforms and readily interchangeable with historic data, is of vital importance as the global community seeks to integrate sequencing data from multiple sources to enable more accurate interpretation of genomic information in the clinic.

A fundamental difference in pre-NGS and NGS variant annotation is in the selection of the gene transcript against which to annotate if a variant is present. For pre-NGS methods a RefSeq transcript is typically used. This often corresponds to an mRNA sequence, usually from a single individual, and may have undergone curation to include the major alleles in a given population [7]. For NGS data, variant detection is made through comparison with the reference human genome sequence, which was generated from several individuals and generally has not been altered to reflect the major alleles in a specific population [8]. This difference can impact variant calling if the RefSeq transcript differs from the reference genome sequence. The *BRCA2* gene exemplifies this issue. The RefSeq transcript NM_000059.3, which has historically been used for pre-NGS *BRCA2* clinical sequencing annotation, has ‘C’ as nucleotide 7397, whereas the reference genome has a ‘T’ at this position, with the corresponding amino acids being alanine and valine respectively. Thus an individual with a ‘C’ at this position would have no variant detected at all in Sanger sequencing data but the same individual would have a nonsynonymous variant c.7397T>C_p.Val2466Ala called in NGS data.

A second important difference is in the description of insertions and deletions (collectively termed ‘indels’). Annotation of indels in Sanger data is undertaken directly in relation to the coding transcript and described in line with the HGVS guidelines which require a variant to be called at the most 3’ position in the coding transcript [3]. In NGS data variant calls are usually reported in a standardized Variant Call Format (VCF), which represents indels at the most 5’ position on the forward strand of DNA; a process called ‘left alignment’ [9]. Adherence to the VCF is not universal; for example, the widely used mpileup command in samtools can report right aligned coordinates [10]. Most existing NGS annotation tools directly annotate the supplied file regardless of left or right alignment [4-6]. These tools thus generate indel calls that are internally inconsistent and externally incompatible because ∼50% of coding transcripts are on the forward DNA strand and ∼50% are on the reverse DNA strand (a small number of genes have overlapping coding transcripts on both strands). Most current NGS annotation tools follow the left aligned input VCF coordinates which position an indel at the most 3’ position if the coding transcript is on the reverse strand (e.g. *BRCA1*), but at the most 5’ position if the coding transcript is on the forward strand (e.g. *BRCA2*).

A further issue is that many indels have different possible representations. Typically, this occurs when the indel occurs in a repetitive region. For example, if a deletion of an ‘A’ is within a polyA tract such as ‘AAAAAA’ it is not possible to definitively know which ‘A’ has been deleted. For some indels these alternative representations have different predicted impacts on the protein and neither pre-NGS nor NGS variant annotation systems currently signpost this important scenario. For example, an indel at the intron-exon boundary could be classified as intronic or exonic depending on which representation is used, with potential significant impact on clinical interpretation (Figure 1).

**Figure 1.**
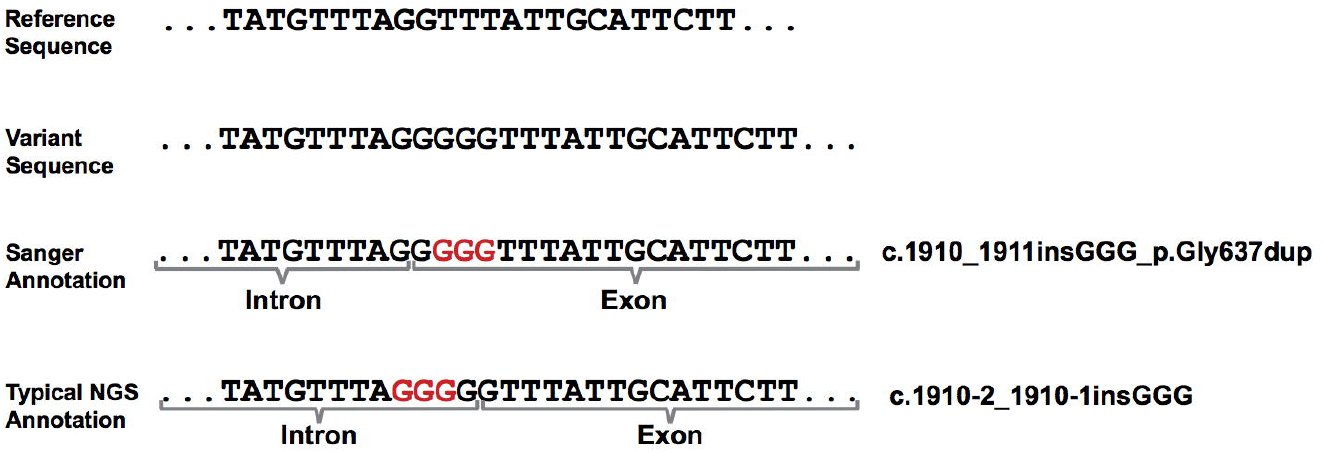
Example of indel with alternative representations. The variant is a ‘GGG’ insertion that overlaps the 5’ boundary of *BRCA2* exon 11. This would be annotated as an inframe glycine duplication in the most 3’ representation, as is standard for clinical annotations, but as an intronic insertion with no impact on coding sequence if left aligned, as is typical for most NGS annotation tools.

These issues became apparent to us through the Mainstreaming Cancer Genetics programme which is using NGS to deliver large-scale, high-throughput, clinical testing of cancer predisposition genes (www.mcgprogramme.com)[11, 12]. Here we describe the tools we developed to address these challenges which we believe have broad relevance and utility for clinical genomics.

## METHODS

### Clinical Sequencing Nomenclature (CSN)

We developed a standardized clinical sequencing nomenclature (CSN) for DNA sequence variant annotation. The aims of CSN are a) to provide a fixed, standardized system in which every variant has a single notation, b) to be identical for all mutation detection methods, c) to use a logical terminology understandable to non-experts, and d) to provide a nomenclature that allows easy visual discrimination between the major classes of variant in clinical genomics. The CSN follows the principles of the HGVS nomenclature, with some minor amendments to ensure compatibility and integration with historical clinical data, whilst also allowing high-throughput automated output from NGS platforms. The CSN is fully detailed in Supplementary Appendix 1.

### Clinical Annotation of VAriants (CAVA)

To provide CSN annotation in a robust and automated fashion, we developed a tool called CAVA (Clinical Annotation of VAriants) which is written in Python. CAVA is DNA ‘strand-aware’, performing coding transcript-dependent alignment so all indels are consistently reported at the most 3’ position in the coding transcript, in line with the HGVS recommendation. CAVA also classifies variants based on their impact on the protein according to a simple ontology (Table 1). Within the CAVA classification system each variant is assigned to a single class to ensure consistency. To facilitate data utilization and comparison with other datasets the Sequence Ontology (SO) classes are also given [13]. CAVA further provides an impact flag which stratifies variants into categories according to predicted severity of impact on protein function, with three default classes: category 1 = ESS, FS, SG, category 2 = NSY, SS5, IF, IM, SL, EE and category 3 = SY, SS, INT, 5PU, 3PU.

**Table 1.** CAVA variant classification system.

| Class | Description |
| --- | --- |
| SG | Stop-gain variant caused by base substitution. |
| ESS | Any variant that alters essential splice-site base (+1, +2, -1, -2). |
| SS5 | Any variant that alters +5 splice-site base but not an ESS base. |
| SS | Any variant that alters splice-site base within the first 8 intronic bases flanking exon (i.e. +8 to -8) but not an ESS or SS5 base. |
| EE | Variant that alters the first or last 3 bases of an exon (i.e. the exon end), but not the frame of the coding sequence. |
| FS | Frameshifting insertion and/or deletion. It alters length and frame of coding sequence. |
| IM | Variant that alters initiating methionine start codon. |
| SL | Variant that causes a stop-loss (i.e. the stop codon is altered). |
| IF | Inframe insertion and/or deletion. It alters length but not frame of coding sequence. |
| NSY | Nonsynonymous variant. It alters amino acid(s) but not coding sequence length. |
| SY | Synonymous variant. It does not alter amino acid or coding sequence length. |
| INT | Any variant in an intron that does not alter splice-site bases. |
| 5PU | Any variant in 5' untranslated region. |
| 3PU | Any variant in 3' untranslated region. |

**Notes:**
A variant can only have one CAVA class. If a variant could potentially be included in more than one class the first class in the list is assigned. For example, a frameshifting deletion that alters the start codon would be CAVA class FS (not IM).
Nonsynonymous is also known as missense. Stop-gain is also known as nonsense.

Default variant annotations outputted by CAVA include the CSN call, variant type (substitution, insertion, deletion or complex), HGNC symbol(s) of affected gene(s), Ensembl transcript identifier(s), within-transcript location(s) (i.e. the exon/intron number or 5’/3’ UTR), the CAVA class, the SO term, the impact category, and the alternate most 5’ annotation (where appropriate). A SNP database can also be used to assign dbSNP identifiers [2].

The user can specify the set of Ensembl transcripts used for variant annotation instead of, or in addition to a default whole exome canonical transcript set provided on installation. CAVA supports overlapping Ensembl transcripts, i.e. a single variant call can be annotated according to multiple transcripts. CAVA also provides various filtering options including removing intergenic variant calls, i.e. calls not overlapping with any included transcripts, or only outputting calls affecting specific genes or genomic regions.

CAVA is lightweight and is easily added to NGS pipelines as it reads variants from VCF files and outputs either a VCF with annotations appended to the original input or an easily parsable tab-separated text file, and both can be written to the standard output. Processing speed can be further increased by parallelization as each line in the VCF file is processed independently. CAVA is fully detailed in Supplementary Appendix 2. CAVA is freely available and can be downloaded from http://www.well.ox.ac.uk/cava

### CAVA exome data annotation

The Exome Aggregation Consortium (ExAC) is a collaborative effort to reanalyze germline exome sequencing data from 61,486 unrelated individuals contributed by a number of disease-specific and population genetic studies [14]. The VCF file containing 10,313,034 variants in release 0.2 was downloaded and annotated by CAVA using a single core.

In-house exome sequencing data was available from 1,000 individuals obtained from the 1958 Birth Cohort Collection (the ICR1000 UK exome series). We used the Illumina TruSeq Exome and sequencing was performed with an Illumina HiSeq2000 generating 2x101bp reads. Reads were mapped to hg19 using Stampy [15] and duplicate reads were flagged with Picard (http://picard.sourceforge.net). Variants were called with Platypus [16], generating raw VCF files. The ICR1000 UK exome data are available from European Genome-phenome Archive [17]. Annotation of the 1,000 VCF files was performed by CAVA in five independent jobs. Each job utilized 15 of the 16 available cores to process files in batches of 15 in parallel with one core per file. Four jobs processed 195 files each, and the fifth processed the remaining 220 files.

### CAVA indel annotation

To evaluate CAVA indel annotation in a typical clinical scenario we used the raw VCF data from a single individual from the ICR1000 series. We excluded intergenic variants and those which only affected intronic or UTR sequence (CAVA classes INT, 3PU, or 5PU).

### CAVA clinical sequence data analysis

We used data from a clinical gene testing laboratory, TGLclinical (www.tglclinical.com), from 25 individuals with *BRCA1* mutations and 25 individuals with *BRCA2* mutations. The mutations had been identified by NGS using the Illumina TruSight Cancer Panel (TSCP) and each mutation was then verified by Sanger sequencing and the Sanger data was used to generate the clinical report. NGS analysis of TSCP used Stampy for alignment [15] and Platypus for variant calling [16]. The default VCF file output from Platypus was used as input for CAVA (v.1.0), VEP (v.77) ANNOVAR (v.2014Jul14) and SnpEff (v.4.0) which were the most recent versions available in November 2014 when the analysis was performed.

## RESULTS AND DISCUSSION

### Clinical Sequencing Nomenclature (CSN)

The CSN is based on the HGVS guidelines to facilitate integration with data generated by pre-NGS methods whilst providing standardization and compatibility with large-scale automated NGS data calling. The full details of the CSN are provided in Supplementary Appendix 1. Key details are outlined here.

CSN provides a single variant call incorporating both the nucleotide and amino acid change (where appropriate), linked by an underscore ‘_’. Currently, most annotation systems provide the nucleotide and amino acid impact separately, either unlinked or variably linked e.g. with semi-colons, commas or a space. This inconsistency causes confusion and impedes data consolidation.

CSN standardizes the description of base substitutions within genes that result in stop-gain (nonsense), nonsyonymous (missense) and synonymous (silent) variants, in a systematic format that allows easy visual discrimination between the classes. This is very helpful in clinical genomics as the variant class is typically not recorded in medical records (Table 2). Historically, HGVS has permitted different notations for stop-gain variants, including ‘X’, ‘*’ and ‘ter’. It is clearly essential that only one notation is used. ‘*’ is not acceptable as this denotes a wild-card in many applications. In the CSN we selected ‘X’. We believe this is preferable to ‘ter’ for three reasons. First, it allows stop-gain variants to be readily discriminated from variants in other classes (Table 2). Second, ‘ter’ is often assumed to denote a specific amino acid, rather than any stop codon, potentially leading to misinterpretation as nonsynonymous. Third, ‘X’ is a very widely used and well-recognised notation for a stop codon in clinical genomics and the scientific literature.

**Table 2.** Comparison of CSN and current nomenclature for exonic base substitutions.

| CSN | Current nomenclature<sup>a</sup> |  |
| --- | --- | --- |
|  | <u>Nucleotide</u> | <u>Amino Acid</u> |
| c.1040A>G_p.Gln347Arg | c.1040A>G | p.Gln347Arg |
| c.1911T>C_p.= | c.1911T>C | p.Gly637Gly |
| c.3264T>C_p.= | c.3264T>C | p.Pro1088Pro |
| c.3515C>T_p.Ser1172Leu | c.3515C>T | p.Ser1172Leu |
| c.3516G>A_p.= | c.3516G>A | p.Ser1172Ser |
| c.5682C>G_p.Tyr1894X | c.5682C>G | p.Tyr1894Ter |
| c.5855T>A_p.Leu1952X | c.5855T>A | p.Leu1952Ter |
| c.6131G>T_p.Gly2044Val | c.6131G>T | p.Gly2044Val |
| c.6675A>G_p.= | c.6675A>G | p.Thr2225Thr |
| c.7558C>T_p.Arg2520X | c.7558C>T | p.Arg2520Ter |
| c.8182G>A_p.Val2728Ile | c.8182G>A | p.Val2728Ile |
| c.9976A>T_p.Lys3326X | c.9976A>T | p.Lys3326Ter |

CSN allows easy visual discrimination between the different classes of exonic base substitutions with '=' denoting a synonymous variant, 'X' denoting a stop-gain variant and the three letter code of the new amino acid denoting a nonsynonymous variant. CSN includes both the nucleotide and amino acid level descriptions to give a single, unique identifier for each variant.
<sup>a</sup>The current nomenclature given is one of several different notation systems currently in use.

For nonsynonymous variants, some annotation systems use a three letter code for amino acids (e.g. p.Gln347Arg), whereas others use a single letter code (e.g. p.Q347R). CSN follows the HGVS preferred recommendation of using the three letter code, which makes it easier to recognize which amino acids are involved: c.1040A>G_p.Gln347Arg. For synonymous variants, some systems include the amino acid code before and after the variant position to indicate there is no change (e.g. c.1911T>C p.Gly637Gly). However, this makes nonsynonymous and synonymous variants difficult to distinguish visually (Table 2). CSN follows the HGVS recommendation of using ‘=’ to show that the amino acid remains the same: c.1911T>C_p.=.

CSN thus provides a simple, distinctive system for exonic base substitutions: ‘X’ indicates a stop-gain variant, ‘=’ indicates a synonymous variant and a three letter code indicates a nonsynonymous variant (Table 2).

Frameshifting indel mutations in CSN are described using only the nucleotide change, as is typical in clinical genomics. Many annotation systems include a hypothetical amino acid change, typically providing the first stop-gain that would occur as a result of the frameshift. However, most frameshifting indels cause nmRNA decay; they do not lead to a truncated protein. Therefore this notation will be incorrect for the great majority of indels. The CSN frameshifting indel notation is also shorter and easier to remember and describe: e.g. *BRCA1* c.246delT (CSN) vs *BRCA1* c.246delT p.Val83LeufsTer5 (VEP). This is important clinically, particularly given the prevalence of this variant class in clinical genomics. CSN positions all indels at their most 3’ position in the coding transcript, as recommended by HGVS. Positioning in relation to the forward strand of DNA, as performed by most NGS annotation tools, is unacceptable as it results in annotation inconsistency as described above.

### Clinical Annotation of Variants tool (CAVA)

To provide CSN annotation in a fast, robust, automated fashion, we developed a tool called CAVA (Clinical Annotation of VAriants). CAVA classifies variants based on a simple, explicit, logical ontology focused on clinical requirements, which avoids historical jargon, such as ‘nonsense’ for a stop-gain mutation. The ontology deliberately focuses on the likely clinical impact of variants, for example, explicitly recognizing any variants that alter the first and last codons of an exon as these often result in splicing defects (Table 1). Additionally, in the CAVA classification system each variant has only one class, to ensure consistency in variant classification. However, the Sequence Ontology (SO) classes are also provided to facilitate analyses and interchange with other datasets [13].

CAVA uses Ensembl transcripts to ensure variants called against the reference human genome are annotated correctly. A default database is included but there is also flexibility to use a bespoke, user-generated transcript database. Importantly, CAVA adjusts for the DNA strand of the coding transcript, so that indels are always called at the most 3’ position in the coding transcript, in line with HGVS and CSN. Furthermore, CAVA flags any variant with potential alternative representations, outputting the alternative annotations as well. This is extremely important clinically as it ensures that, where appropriate, the most deleterious potential consequence of a variant can be investigated (e.g. Figure 1). Highlighting variants with alternative possible annotations also facilitates comparisons with variant sets annotated with other tools. Examples of the default CAVA outputs are shown in Table 3.

**Table 3.** Example default output of CAVAv1.0.

| CH R | POS | REF | ALT | QUAL | FILTER | TYPE | ENST | GENE | TRINFO | LOC | CSN | CLASS | SO | IMPACT | ALTANN | ALT CLASS | ALT SO |
| --- | --- | --- | --- | --- | --- | --- | --- | --- | --- | --- | --- | --- | --- | --- | --- | --- | --- |
| 1 | 12009955 | C | T | 200 | PASS | Substitution | ENST00000196061 | PLOD1 | +/-40.8kb/19/2.9kb | Ex3 | c.294C>T_p.= | SY | synonymous_variant | 3 | . | . | . |
| 1 | 12919891 | G | T | 200 | PASS | Substitution | ENST00000240189 | PRAMEF2 | +/-4.8kb/4/1.6kb | Ex3 | c.631G>T_p.Glu211X | SG | stop_gained | 1 | . | . | . |
| 1 | 14106394 | A | ACTC | 200 | PASS | Insertion | ENST00000235372 | PRDM2 | +/-120.2kb/10/7.9kb | Ex8 | c.2107_2109dupCCT_p.Pro703dup | IF | inframe_insertion | 2 | c.2104_2105insCTC_p.Pro703dup | . | . |
| 1 | 15789297 | A | C | 200 | PASS | Substitution | ENST00000359621 | CELA2A | +/-15.4kb/8/0.9kb | Ex4 | c.297A>C_p.= | SY | synonymous_variant | 3 | . | . | . |
| 1 | 15812432 | A | G | 200 | PASS | Substitution | ENST00000375910 | CELA2B | +/-15.3kb/8/0.9kb | Ex6 | c.530A>G_p.Gln177Arg | NSY | missense_variant | 2 | . | . | . |
| 1 | 16727305 | G | GCTT | 200 | PASS | Insertion | ENST00000335496 | SPATA21 | - /38.8kb/13/2.0kb | Ex11 | c.1081_1083dupAAG_p.Lys361dup | IF | inframe_insertion | 2 | c.1078_1079insAGA_p.Lys361dup | . | . |
| 1 | 22310824 | T | C | 200 | PASS | Substitution | ENST00000337107 | CELA3B | +/-12.3kb/8/0.9kb | Ex6 | c.642T>C_p.= | EE | splice_region_variant\|synonymous_variant | 2 | . | . | . |
| 1 | 31905889 | A | ACAG | 200 | PASS | Insertion | ENST00000373710 | SERINC2 | +/-25.1kb/11/2.1kb | Ex10 | c.1129_1131dupCAG_p.Gln377dup | IF | inframe_insertion | 2 | c.1116_1117insCAG_p.Gln377dup | . | . |
| 1 | 36937059 | A | G | 200 | PASS | Substitution | ENST00000373103 | CSF3R | - /17.2kb/17/3.5kb | Ex10 | c.1260T>C_p.= | SY | synonymous_variant | 3 | . | . | . |
| 1 | 38023316 | C | T | 200 | PASS | Substitution | ENST00000296218 | DNALI1 | +/-9.9kb/6/2.6kb | Ex2 | c.260C>T_p.Ala87Val | NSY | missense_variant | 2 | . | . | . |
| 1 | 43771016 | TA | T | 200 | PASS | Deletion | ENST00000372476 | TIE1 | +/-22.1kb/23/3.9kb | In3/4 | c.484+5delA | SS5 | splice_donor_5th_base_variant | 2 | c.484+3delA | . | . |
| 1 | 54605319 | G | GC | 200 | PASS | Insertion | ENST00000371330 | CDCP2 | - /14.8kb/4/2.7kb | Ex4 | c.1223_1224insG | FS | frameshift_variant | 1 | . | . | . |
| 1 | 55251689 | T | C | 200 | PASS | Substitution | ENST00000371276 | TTC22 | - /21.6kb/7/3.3kb | Ex5 | c.987A>G_p.= | SY | synonymous_variant | 3 | . | . | . |
| 1 | 55603581 | T | TA | 200 | PASS | Insertion | ENST00000294383 | USP24 | - /149.0kb/68/10.8kb | In26/27 | c.2929-5dupT | SS | intron_variant\|splice_region_variant | 3 | c.2929-9_2929-8insT | INT | intron_variant |
| 1 | 60503762 | T | C | 200 | PASS | Substitution | ENST00000371201 | C1orf87 | - /83.4kb/12/2.0kb | Ex6 | c.765A>G_p.= | SY | synonymous_variant | 3 | . | . | . |
| 1 | 62232031 | C | T | 200 | PASS | Substitution | ENST00000371158 | INADL | +/-421.4kb/43/8.5kb | Ex4 | c.270C>T_p.= | SY | synonymous_variant | 3 | . | . | . |
| 1 | 67155862 | TCTC | T | 200 | PASS | Deletion | ENST00000371037 | SGIP1 | +/-210.8kb/25/4.6kb | In16/17 | c.1444-8_1444-6delCCT | SS | intron_variant\|splice_region_variant | 3 | c.1444-10_1444-8delCTC | . | . |

In addition to providing consistent clinical annotations, CAVA is freely available and designed to be lightweight, flexible and to be easily appended to any NGS pipeline to provide high utility for clinical and research applications. Full details of CAVA are provided in Supplementary Appendix 2.

### CAVA exome annotation

To evaluate performance in annotating large variant datasets we used CAVA to annotate the Exome Aggregation Consortium (ExAC) data. Annotation of 10,313,034 variants took 13.44 hours i.e. at a rate of 14,234 variants / minute. Faster annotation would be easily attainable with parallelization. This annotation was also of practical utility because the ExAC data in r0.2 provides only the amino acid change for exonic base substitutions which impedes clinical utilisation and comparison with other data, particularly since the degeneracy of the genetic code allows different mutations at the nucleotide level to result in the same mutation at the amino acid level.

To evaluate CAVA performance in real-time whole exome annotation we analyzed the ICR1000 UK exome series using parallelized annotation in batches of 15 exomes. The average file had 170,900 variants (range 108,400-225,000), and the 1,000 exomes were annotated in ∼6.5 hours. We used the data from one individual to evaluate CAVA indel annotation in a typical clinical scenario. This individual had 731 different indels, which were distributed equally amongst genes with coding transcripts on the forward and reverse DNA strands (Supplementary Table 1). 92% (675/731) of indels had an alternative representation and would thus be represented differently in left aligned and right aligned data. Annotation tools that do not incorporate the strand of the coding transcript would thus lead to calls discrepant with clinical annotation for 339 indels (those in genes transcribed from the forward DNA strand); 46% of all indels in this individual. Furthermore, 370 indels had an alternative representation that was also of a different class (Supplementary Table 1). This includes 27 indels for which only one representation was predicted to cause premature protein truncation (either FS or ESS). The functional and clinical implications of truncating and non-truncating variants are potentially very different and it is thus essential in clinical genomics that such variants are highlighted.

### CAVA clinical annotation

To evaluate and compare CAVA and standard NGS annotation tools for indels in the clinical setting we used data from a *BRCA1* and *BRCA2* clinical testing laboratory, in which testing is performed by NGS panel analysis with pathogenic indel mutations confirmed by Sanger sequencing. 25 *BRCA1* and 25 *BRCA2* indels were evaluated (Supplementary Table 2). CAVA provided annotations consistent with the clinical report for all 50 mutations. Additionally CAVA flagged that alternate annotations were possible for 36 mutations, though none altered the class (i.e. all possible representations result in a frameshift). By contrast, only 8/25 (32%) of the *BRCA2* indels were correctly clinically annotated by other tools (Supplementary Table 2).

## CONCLUSIONS

We have highlighted in this paper some of the rudimentary problems in variant annotation that are hindering the large-scale implementation of genomic medicine that NGS is poised to deliver. A fundamental problem is the absence of consistent annotation of variants in the clinic. We here introduce the CSN a nomenclature for clinical sequence data which we believe can serve as the foundation of an integrative, cross-platform annotation system optimized for technological, informatic and clinical requirements. There remain several areas requiring standardization, for example a defined, consensus set of gene transcripts against which to perform clinical annotation must be decided. Expansion of CSN to provide standardization of annotation of additional variant classes such as larger exonic deletions and duplications will also be required. Ongoing CSN iteration, performed by an appropriately representative group, and with all modifications explicitly detailed and versioned, will thus be essential.

We also show the profound impact that the strandedness of transcripts can have on the annotation and interpretation of indels. It is essential that all variant annotation tools recognize and address this issue. We have developed CAVA, a freely available, lightweight annotation tool that can be readily appended to NGS pipelines and which incorporates the transcript strand to provide consistent, clinically appropriate indel calls. Equally importantly, CAVA highlights indels that have possible alternative annotations so that fully-informed clinical interpretation can be performed.

We have implemented CSN using CAVA in a clinical gene testing lab performing cancer predisposition gene panel testing, allowing robust, high-throughput gene testing, adhering to clinical testing standards, to be delivered. The problems we highlight and the solutions we have developed are generic and therefore should have broad relevance and utility in genomic medicine.

## LIST OF ABBREVIATIONS

CSN: Clinical sequencing nomenclature
ExAC: Exome aggregation consortium
HGNC: HUGO gene nomenclature committee
HGVS: Human genome variation society
SO: Sequence ontology
TSCP: Illumina TruSight cancer panel
UTR: Untranslated region
VCF: Variant call format

## URL

1958 Birth Cohort details available at http://www2.le.ac.uk/projects/birthcohort

CAVA is available from http://www.well.ox.ac.uk/cava

dbSNP information for rs80357713 http://www.ncbi.nlm.nih.gov/SNP/

Samtools is documented at http://www.htslib.org/

Exome Aggregation Consortium (ExAC), Cambridge, MA (URL: http://exac.broadinstitute.org)

The ICR1000 series [Accession number: EGAS00001000971] from European Genome-phenome Archive https://www.ebi.ac.uk/ega/datasets/EGAD00001001021

Illumina TruSight Cancer Panel (TSCP) http://www.illumina.com/products/trusight_cancer.html

## COMPETING INTERESTS

The authors declare no conflict of interest.

## AUTHORS’ CONTRIBUTIONS

NR, ERu and AR developed CSN. MM, ERu, GL and NR participated in the design of CAVA, the code for which was written by MM. ERu performed the ICR1000 and clinical data evaluations. MC performed the ExAC analysis. AR and ERa generated the ICR1000 data. VC, SS, SM, AS provided the clinical laboratory sequence data. The manuscript was written by NR, ERu, MM and AS. All authors read and approved the final manuscript.

## SUPPLEMENTARY MATERIALS

**Supplementary Table 1**. Indels in the exome of an individual in the ICR1000 series

**Supplementary Table 2**. Comparison of Clinical (Sanger) and NGS annotation of *BRCA1* and *BRCA2*

mutations

**Supplementary Appendix 1**. Clinical Sequencing Nomenclature (CSN) description

**Supplementary Appendix 2.** Clinical Annotation of Variants (CAVA) description

## ACKNOWLEDGEMENTS

We thank Andrew Rimmer, Fran Smith and Razvan Sultana for helpful contributions. We are grateful to Sandra Hanks, Emma Ramsay, Silvana Powell, Imran Hussain and Ann Strydom for management of the genetics laboratory resources used.

This work made use of samples generated by the 1958 Birth Cohort (NCDS). Access to these resources was enabled via the 58READIE Project funded by Wellcome Trust and Medical Research Council (grant numbers WT095219MA and G1001799). A full list of the financial, institutional and personal contributions to the development of the 1958 Birth Cohort Biomedical resource is available at http://www2.le.ac.uk/projects/birthcohort.

The authors would like to thank the Exome Aggregation Consortium and the groups that provided exome variant data for comparison. A full list of contributing groups can be found at http://exac.broadinstitute.org/about.

Rahman acknowledges support from the NIHR RM/ICR Biomedical Research Centre.

*Funding*: This work was supported by Wellcome Trust Award 098518/Z/12/Z.

